# Simultaneous mesoscopic and two-photon imaging of neuronal activity in cortical circuits

**DOI:** 10.1101/468348

**Authors:** D Barson, AS Hamodi, X Shen, G Lur, RT Constable, JA Cardin, MC Crair, MJ Higley

## Abstract

Spontaneous and sensory-evoked activity propagates across spatial scales in the mammalian cortex but technical challenges have generally precluded establishing conceptual links between the function of local circuits of neurons and brain-wide network dynamics. To solve this problem, we developed a method for simultaneous cellular-resolution two-photon calcium imaging of a local microcircuit and mesoscopic widefield calcium imaging of the entire cortical mantle in awake, behaving mice. Our method employs an orthogonal axis design whereby the mesoscopic objective is oriented downward directly above the brain and the two-photon objective is oriented horizontally, with imaging performed through a glass right angle microprism implanted in the skull. In support of this method, we introduce a suite of analysis tools for relating the activity of individual cells to distal cortical areas, as well as a viral method for robust and widespread gene delivery in the juvenile mouse brain. We use these methods to characterize the diversity of associations of individual, genetically-defined neurons with cortex-wide network motifs.

## Introduction

In the mammalian neocortex, single neurons integrate excitatory and inhibitory synaptic inputs arising from both local circuits and long-range projections originating in various cortical and sub-cortical structures^1–3^. This connectivity gives rise to functional networks distributed throughout the cortex, thought to be dedicated to processing various streams of information relevant for cognitive abilities, including sensory and motor representations^4,5^. Interestingly, anatomical, electrophysiological, and imaging studies have demonstrated distinct local and large-scale connectivity associated with varied feature encoding even for neighboring neurons in a single region^6–10^. Nevertheless, most experimental protocols are confined to single areas or modalities, limiting the ability to link the function of local circuits to more global cortical dynamics. The challenge stems largely from methodological hurdles to simultaneous measurement of neural activity across wide spatiotemporal scales. For example, electrophysiological recordings and 2-photon calcium imaging have been highly successful in monitoring the output of single neurons. In contrast, electroencephalography, fMRI, and more recently, mesoscopic calcium imaging, have been successful at reporting broad levels of activity across the cortex. Recent studies have sought to bridge this gap by expanding the capabilities of existing techniques^11–13^, but methods for relating cortical function across these scales remain elusive.

Here, we introduce a novel approach for performing simultaneous measurements of the micro-scale activity of single neurons and the meso-scale activity of areas across the cortical mantle by combining two-photon and widefield calcium imaging in a new imaging system. We also develop an analytical framework for relating activity measured with these two modalities. Widefield calcium imaging is performed through an intact-skull preparation while two-photon imaging is performed using a long working-distance objective directed orthogonally to the brain surface through a glass microprism implanted above the cortex. Compared to previously described methods pairing extra-cellular electrophysiology with mesoscopic calcium imaging^14,15^, our method has several advantages. With two-photon imaging, we can record from hundreds of neurons simultaneously, targeting genetically-defined (often sparse) cell populations. Furthermore, we can easily follow the same cells over days or weeks, allowing us to monitor the stability and flexibility of cortical circuits.

Here, we apply our simultaneous imaging method to transgenic mice expressing GCaMP6 in all cortical pyramidal neurons. We provide the first direct assessment of the cellular underpinnings of the mesoscopic calcium signal, finding that it is highly correlated with the local neuropil. We also describe the diversity of cortex-wide network membership for individual cells in the primary somatosensory cortex (S1) of awake, juvenile mice. Using a novel functional parcellation for widefield calcium imaging data^16^, we find that activity-based segmentation of cortical networks is more effective than a commonly used anatomically-based approach. Finally, we leverage the cell-type specificity afforded by the use of genetically-encoded indicators to reveal the association of vasoactive intestinal peptide-expressing interneurons (VIP-INs) with distal cortical areas across behavioral state.

## Results

### Design of a dual-axis microscope for simultaneous 2-photon and mesoscopic imaging

We developed a method to simultaneously record the single cell-resolution activity of hundreds of neurons within a cortical area and the mesoscopic activity of all other areas across the cortical mantle in both developing and adult mice. We employ a “dual-axis” design^17^ that combines a widefield epifluorescence “mesoscope” using an objective positioned normal to the surface of the animal’s skull with a two-photon microscope using an ultra-long working distance (20 mm) objective positioned tangential to the skull surface and orthogonal to the mesoscope objective (Figure 1a). In order to reflect the two-photon excitation and emission paths to/from the microscope, we implant a square right angle glass microprism with an uncoated hypotenuse into a small craniotomy where it is placed on the brain surface. The uncoated microprism enables imaging the same brain tissue with either the two-photon or mesoscopic system (Figure 2b).

**Fig. 1.**
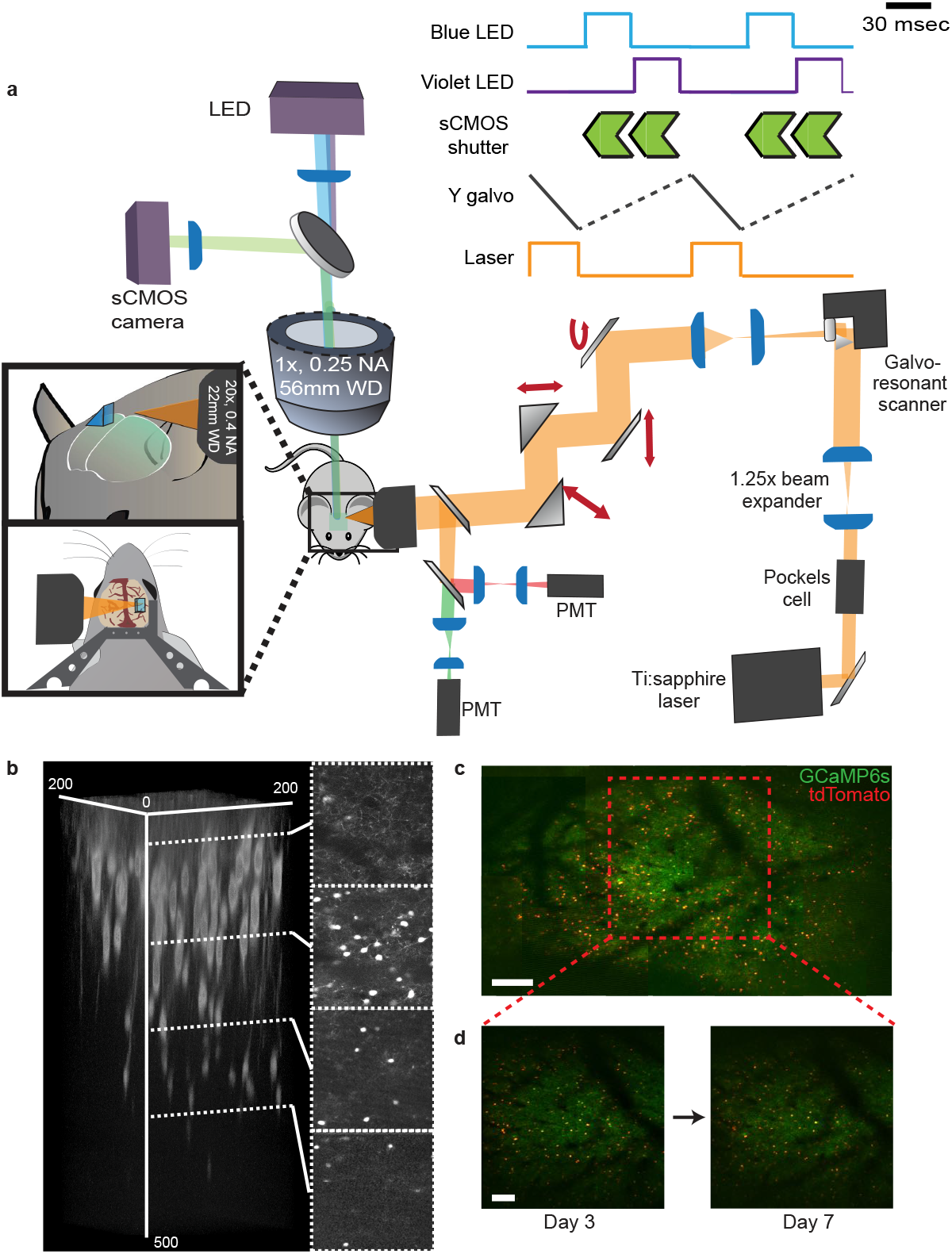
Design of a dual-axis microscope for simultaneous widefield and two-photon imaging. **a**, Schematic overview of the dual-axis microscope. Left insets show the position of the two-photon objective relative to an implanted glass microprism and design of a custom headpost. Right inset shows timing of the widefield LED illumination, widefield sCMOS detector, two-photon excitation laser, and two-photon galvo scanner (proxy for acquisition time). **b**, Three-dimensional reconstruction of a Z-stack of images acquired with the two-photon microscope through a glass microprism with representative images from indicated depths to the right. Images were acquired in an awake mouse expressing the red fluorophore tdTomato in VIP-INs. Indicated dimensions are in microns. **c**, Stitched image of tiled two-photon acquisitions across the entire field-of-view accessible through a 2 mm glass microprism in a mouse expressing pan-neuronal GCaMP and tdTomato in VIP interneurons. Scale bar is 200 µm. **d**, Same field-of-view, corresponding to the red box in c, demonstrating repeated imaging of the same cells on day 3 and day 7 after microprism implantation. Scale bar is 100 µm

**Fig. 2.**
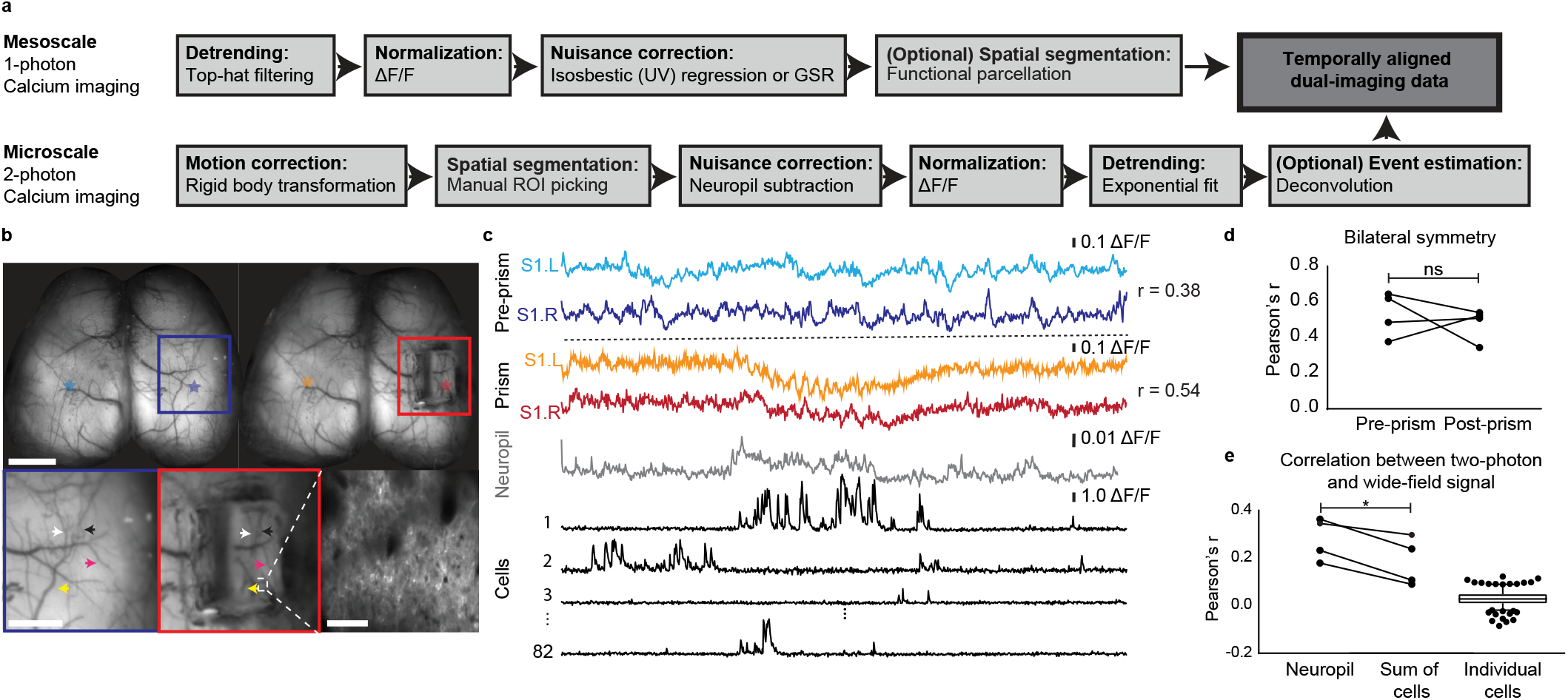
Analysis of simultaneously acquired micro- and mesoscale calcium imaging data in transgenic mice. **a**, Overview of the analysis pipeline for pre-processing and combining simultaneously acquired widefield and two-photon calcium imaging data. **b**, Top: Widefield images of the same animal acquired before and after microprism implantation. Colored stars correspond to regions-of-interest for traces in c. Darker colors are the same area in which the two-photon acquisition was performed. Scale bar is 2 mm. Bottom left, middle: Expanded images corresponding to colored boxes in top images. Colored arrowheads highlight matching blood vessels on the surface of the brain before and after microprism implantation. Scale bar is 1 mm. Bottom right: two-photon field-of-view corresponding to dark colored stars in the top images. Scale bar is 50 µm. **c**, Fluorescence calcium imaging traces (ΔF/F) corresponding to regions-of-interest indicated in b. Prism, neuropil, and cell traces were acquired simultaneously, whereas pre-prism traces were acquired during a previous imaging session. R values between widefield traces are Pearson’s correlation. **d**, Pearson’s correlation between bilateral S1 widefield pixels before and after prism implantation. P = 0.5996, paired two-tailed t test. **e**, Pearson’s correlation between mean fluorescence of widefield pixels corresponding to the two-photon field-of-view and mean fluorescence of all neuropil pixels, all cell pixels, or individual cell pixels in the two-photon field-of-view. P = 0.0133, two-tailed t test, n = 4 trials across 3 animals. Box-and-whisker plot of cell correlations shows median and 5-95 percentile values.

In order to prevent optical cross-talk between the two imaging systems, we interleave the acquisition of widefield epifluorescence and two-photon frames. Triggered LED illumination can rapidly drive alternating widefield excitation (see below). Emitted signals are collected with a high quantum efficiency sCMOS camera. A Pockels cell similarly modulates the intensity of the titanium-sapphire laser source for two-photon excitation (Figure 1a), and emitted photons are collected by photomultiplier tubes. Thus, the start of each widefield frame lags the start of each two-photon frame by 34 – 67 milliseconds, comparable to the rise time and substantially less than the decay time of GCaMP6f for a single action potential^18^, and is therefore effectively simultaneous. For experiments where calcium-dependent and -independent widefield signals are collected, a frame of violet (395 nm) illumination is interleaved with a frame of cyan (470 nm) illumination yielding an overall acquisition rate of up to 15 Hz. Notably, two-photon imaging through the microprism preserves the ability to image cells up to 400 microns below the cortical surface (Figure 1b) underneath the 2 mm^2^ prism (Figure 1c). Moreover, the same cells can be imaged across multiple days (Figure 1d).

### Simultaneous micro- and meso-scale calcium imaging in transgenic mice expressing GCaMP6 in cortical pyramidal neurons

We developed an analysis pipeline to facilitate comparison of cellular-resolution and meso-scale calcium imaging, leveraging our experience with these two techniques individually (Figure 2a). Specifically, we motion correct our two-photon data by rigid body transformation^19^ and perform region-of-interest (ROI) selection and neuropil subtraction by standard methods^9,18^. To eliminate minor photobleaching, we detrend all images using top-hat filtering and fitting the baseline of each ROI signal to a second-order exponential. We also exclude any cell for which the skewness of its signal is less than 0.5 (see Methods). For the mesoscopic data, we perform top-hat filtering to remove slow fluctuations over time and regress out the violet-illuminated isosbestic GCaMP6 signal (or the global signal for data collected without interleaved violet/cyan illumination) to remove fluorescence changes attributable to motion and hemodynamic artefacts^20^.

To test whether mesoscale activity patterns change as a result of the dual-imaging preparation, we acquired mesoscopic calcium data in a cohort of adult transgenic mice expressing GCaMP6f in cortical pyramidal neurons (Slc17a7-cre/Camk2a-tTA/TITL-GCaMP6f, P60-70, n = 3) before and after implantation of the glass microprism over right S1. After prism implantation, much of the area under the microprism continues to be optically accessible to the mesoscope, (arrowheads indicate matching blood vessels in Figure 2b). In awake mice, the correlation of spontaneous activity between right and left S1 is typically high. Comparing the bilateral correlation between right and left S1 before and after prism implantation, we also observe high correlation values and no difference resulting from prism implantation (r = 0.534 +/- 0.063 before, r = 0.4809 +/- 0.046 after, p = 0.6, Figure 2c,d), suggesting that the prism does not does not significantly distort imaging through the mesoscope.

Having simultaneous mesoscopic and cellular-resolution optical access to the same brain area also permits us to directly determine the relationship between the mesoscopic signal and its cellular sources, indirectly inferred in previous studies^15,20,21^. We find that the mesoscopic signal is more highly correlated with layer 2/3 neuropil than the summed activity of all cells (neuropil r = 0.280 +/- 0.044; sum of cells r = 0.183 +/- 0.050, p = 0.013) in the two-photon field-of-view (Figure 2e). Furthermore, individual cells contribute negligibly to the mesoscopic signal (r = 0.006 – 0.052, [25%ile – 75%ile]). Thus, the mesoscopic signal corresponds most directly to axonal and dendritic structures whose cell bodies may not be present in the field of view.

### Simultaneous micro- and meso-scale calcium imaging reveals diverse cortex-wide networks for neighboring pyramidal neurons

We performed simultaneous two-photon imaging of layer 2/3 neurons in mouse S1 and mesoscopic imaging of the entire cortical surface in a cohort of juvenile transgenic mice expressing GCaMP6f in cortical pyramidal neurons (Slc17a7-cre/Camk2a-tTA/TITL-GCaMP6f, P17 – 19, n = 3; Figure 3a,b). Mice were awake and free to run on a circular treadmill^22,23^. Previous work combining extracellular electrophysiology and widefield calcium imaging suggested that S1 neurons in adults participate in a cortex-wide network including bilateral S1 and medial motor cortex, consistent with the known reciprocal connection between these two areas^14,24^. We sought to determine the extent to which this network structure has already emerged in the juvenile cortex, shortly after the closure of the critical period for horizontal connections within layer 2/3 of S1^25^ and the development of callosal connections^26^.

**Fig. 3.**
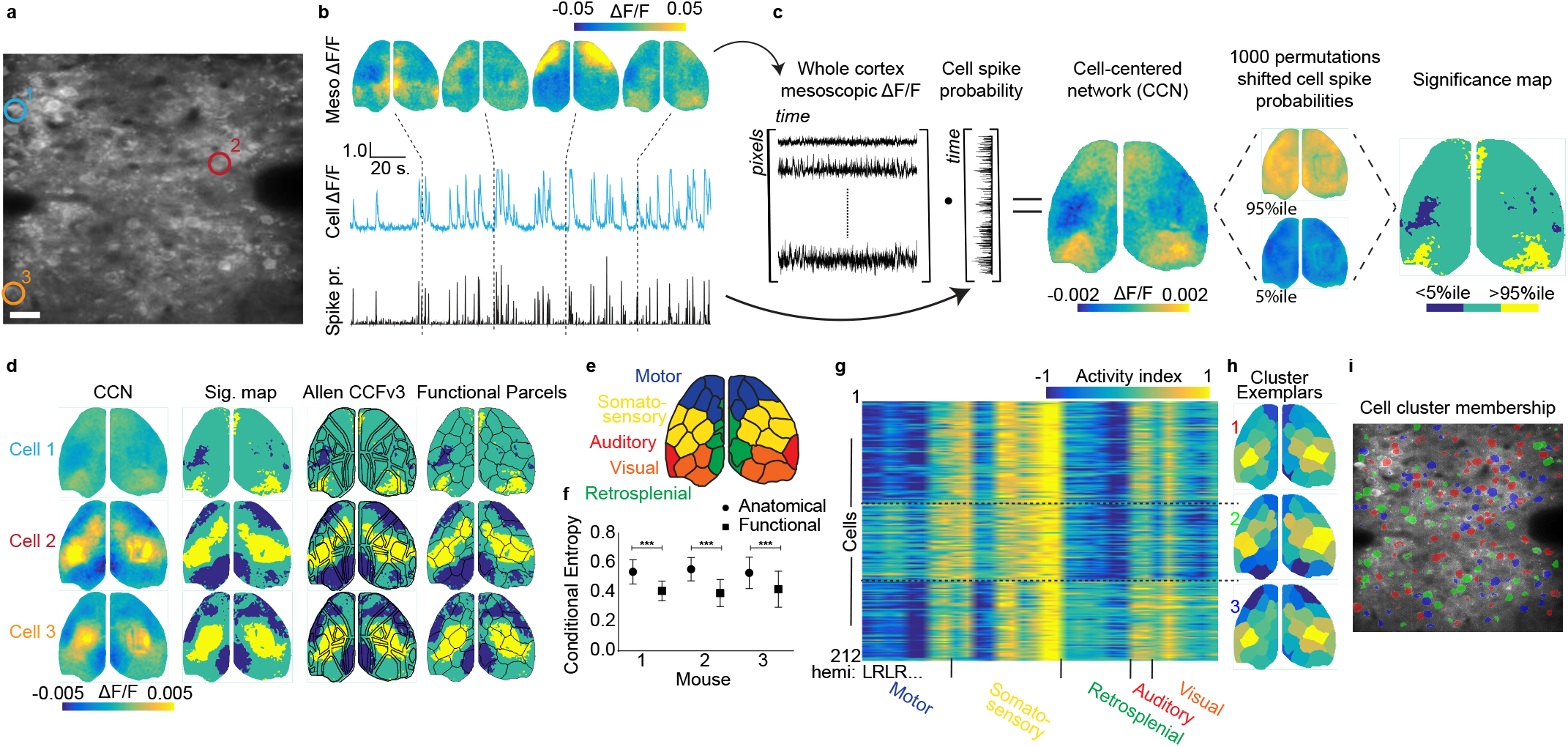
Diverse participation of pyramidal cells in global cortical networks revealed by simultaneous imaging. **a**, Example mean two-photon field-of-view of pyramidal cells in a P17 mouse during simultaneous imaging. Colors of circled cells correspond to colors in b, d. Scale bar is 20 µm. **b**, Example simultaneously acquired widefield ΔF/F images with ΔF/F trace and deconvolved spike probability for a single cell. **c.** Schematized procedure for calculating cell-centered networks (CCNs) and significance maps. **d**, Left: Example CCNs for three cells indicated in a. Middle left: corresponding significance maps. Middle right: significance maps overlaid with an anatomical parcellation based on the Allen CCFv3. Right: significance maps overlaid with a functional parcellation calculated for that mouse. **e**, Illustration of the functional parcellation with regions labelled based on correspondence with the anatomical parcellation. **f**, Conditional entropy of significance maps given the anatomical or functional parcellation for three mice. Lower values indicate better fit. P < 0.001, paired two-tailed t-test for each mouse, n = 212, 43, 33 significance maps. **g**, Activity index calculated from all significance maps for a single animal using the functional parcellation. Higher values indicate a large number of pixels that are significantly co-active with each cell. Cells are clustered into three groups (see methods). **h**, Exemplars from the three clusters in g with parcels colored by their activity index. **I**, Two-photon field-of-view, same as in a, with pixels colored to indicate membership of individual cells in the three clusters shown in g, h.

To relate the activity of each imaged cell to all other simultaneously imaged areas across the cortical mantle, we deconvolved the cell ΔF/F^27^ to estimate the relative probability of cell spiking (Figure 3b). We then computed the dot product with the ΔF/F for each widefield pixel, producing an activity-weighted map illustrating the functional connectivity of that neuron with distal cortical regions. To determine which pixels in these cell-centered networks (CCNs) are significantly activated or deactivated, we compared each CCN to a null distribution generated by 1000 random shuffles of the timing of cell activity (Figure 3c). We assess the reliability of CCNs by splitting each trial in half and computing network structure for the first and second half of the trial. Note that cells with a split-half Pearson's correlation less than 0.3 were excluded from further analysis. Three example significance maps demonstrate that neighboring S1 neurons are coupled with distinct (although overlapping) distal cortical regions (Figure 3d).

To quantify the diversity of CCNs, we superimposed a 16 node-per-hemisphere anatomical parcellation based on the Allen CCFv3 atlas^6^. However, we found that these parcels poorly fit the contours of the CCNs (Figure 3d), likely resulting from a combination of individual differences in the functional organization of spontaneous activity across mice, as well as subtle differences in the angle of each mouse’s skull relative to the wide-field objective. Therefore, we utilized a multi-graph k-way spectral clustering algorithm^28^ to determine a functional 16 node-per-hemisphere parcellation for each mouse based on awake, spontaneous mesoscopic calcium imaging data recorded prior to prism implantation (Figure 3e). Remarkably, this segmentation more closely approximated the contours of activations in the CCNs (Figure 3d). To compare the quality of fit provided by the two parcellations, we calculated the conditional entropy of each CCN given the parcellation (anatomical or functional). Lower conditional entropy indicates a better fit. We found that across all mice, the functional parcellation significantly outperformed the anatomical parcellation (Figure 3f, p < 0.001).

For each cell, we calculated an activity index vector, which is the number of activated and deactivated pixels in each parcel normalized by the number of pixels in the parcel (rows in Figure 3g) and used it to cluster the cells to identify the three most prominent mesoscale patterns of activity coincident with the activity of recorded cells in each mouse (Figure 3h, Supplementary Figure 1). For all the three mice in our sample, the most prominent patterns typically included S1 activation bilaterally but were otherwise varied, with some showing activation of medial motor cortex (exemplar 2), deactivation of lateral motor cortex (exemplar 3), and deactivation of visual (exemplar 2) and retrosplenial cortex (exemplar 1 and 2) coincident with S1 cell activity. Cells belonging to each cluster are intermixed within the two-photon field-of-view (Figure 3i). This greater diversity of cortex-wide network membership than previously reported^14^ may be attributable to either the immature developmental stage of the mice used in our study or the greater number of cells assessed.

### Viral vector-driven whole-brain expression of GCaMP6

Transgenic expression of GCaMP6 in cortical neurons is limited by the complexity of interbreeding various reporter lines, often necessitating the combination of several strains to drive conditional expression in targeted cell types^29,30^. Here, we optimized a viral approach to achieve widespread and robust expression of GCaMP6 throughout the mouse brain using serotype 9 adeno-associated virus (AAV9) injection into the transverse sinuses of early postnatal animals.

AAV9 crosses the blood-brain barrier^31^, and recently published work has demonstrated whole-brain gene delivery by retro-orbital injection of this serotype as well as other engineered AAV variants^32^. While retro-orbital injections are difficult and disruptive in early postnatal mice due to eyelid closure, the transverse sinuses are easily accessible along the posterior edge of the cortex.

We reasoned that injection of AAV9 directly into the venous circulation of the brain would have a high likelihood of successful gene delivery. Therefore, we performed sinus injections of AAV9-Syn-GCaMP6s at P1 (Figure 4a) in wild-type mice and observed widespread expression of green fluorescence as early as P14 (Figure 4b,c). Using this method, we labeled 48.3 +/- 4.2% and 46.4 +/- 15.8% of cortical neurons at P14 and P21, respectively (Figure 4d,e; P14, n = 3; P21, n = 3), and 65.8 +/- 9.3% and 40.8 +/- 9.5% of thalamic neurons (Figure 4f,g; P14, n = 3; P21, n =2). Importantly, this method resulted in sufficiently bright GCaMP6 expression for use with *in vivo* imaging at these ages (Figure 4h,i).

**Fig. 4.**
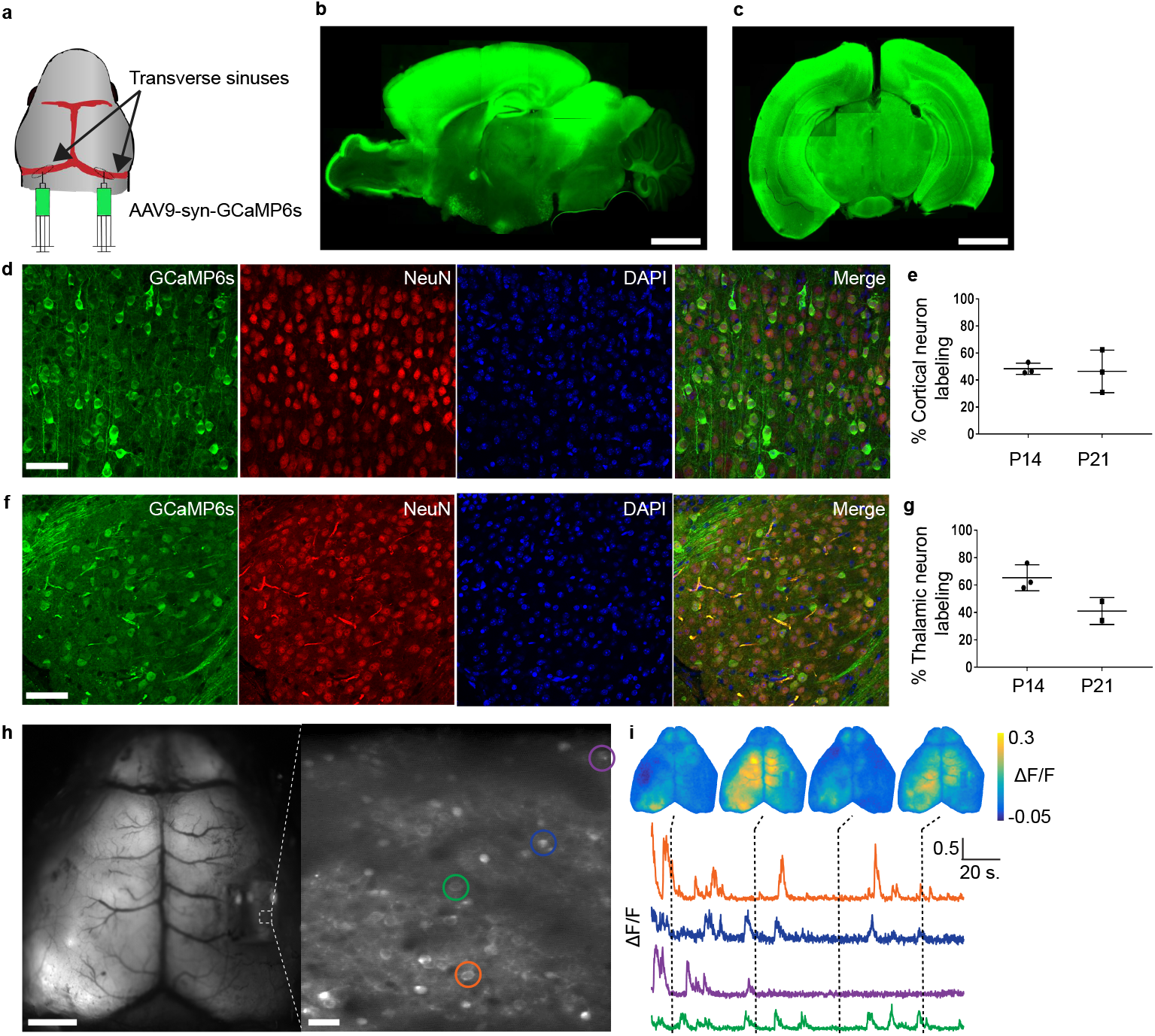
Injection of AAV9 into the transverse sinuses of neonatal mice produces robust and widespread GCaMP expression. **a**, Schematic showing sites of viral injection at P1. **b**, Example sagittal section of a P21 mouse brain showing widespread and even expression of GCaMP6s across the cortex and several other brain regions including hippocampus, midbrain, and thalamus. Scale bar is 2 mm. **c**, Example coronal section. Scale bar is 2 mm. **d**, Confocal images showing GCaMP6s expression in mouse cortex at P14. Left: GCaMP6s, middle left: NeuN, middle right: DAPI, right: merge. Scale bar is 40 μm. **e**, Quantification of cortical neuron labeling at P14 and P21. Each data point on the plot represents an individual brain. **f**, Same as d, but for thalamus. **g**, Same as e, but for thalamus. **h**, Mean images from simultaneously acquired widefield (left, scale bar is 2 mm) and two-photon (right, scale bar is 20 μm) imaging. **i**, Example simultaneously acquired widefield ΔF/F images and cellular ΔF/F traces from cells indicated in h.

### VIP-INs belong to a single large-scale network associated with arousal

As noted above, one advantage to using viral vectors for pan-cortical GCaMP6 expression is the possibility of combining functional imaging with genetic labeling of sparse neuronal populations not easily accessible with electrophysiology. For example, VIP-INs account for less than 2% of cortical neurons^33^. However, they have been strongly implicated in the arousal-dependent modulation of cortical activity via disinhibition of local pyramidal cells^34^. Moreover VIPIN activity is critical for the proper development of cortical circuitry^23^, but little is known about their relationship to overall cortical dynamics.

To investigate the coupling of VIP-INs to cortical mesoscopic networks, we performed sinus injections of AAV9-Syn-GCaMP6s at P1 in mice transgenically expressing the red fluorophore tdTomato in VIP-INs (see methods). We also used local cortical injections of AAV5-CAG-FLEX-GCaMP6s into right S1 at P6 to boost the GCaMP6 expression in VIP-INs. At P16, we head-posted and acclimatized the mice to wheel running, followed by prism implantation and dual-imaging between P17-P20. For all mice, we also recorded wheel velocity and whisker movements by videography (Figure 5a-c) as markers of arousal level^22,35^. Across four mice, we imaged a total of 455 red fluorescent VIP-INs, of which 50 cells were included in subsequent analyses (skewness greater than 0.5). Excluded cells represent a mixture of inactive cells and cells that did not express GCaMP. We also recorded 481 presumptive pyramidal (non-tdTomato-expressing) cells, of which 342 were included in subsequent analyses. For each active cell, we calculated the correlation of its ΔF/F with whisker motion energy and running speed and categorized cells as significantly modulated (positively or negatively) nor non-modulated by whisking or running (Figure 5d).

**Fig. 5.**
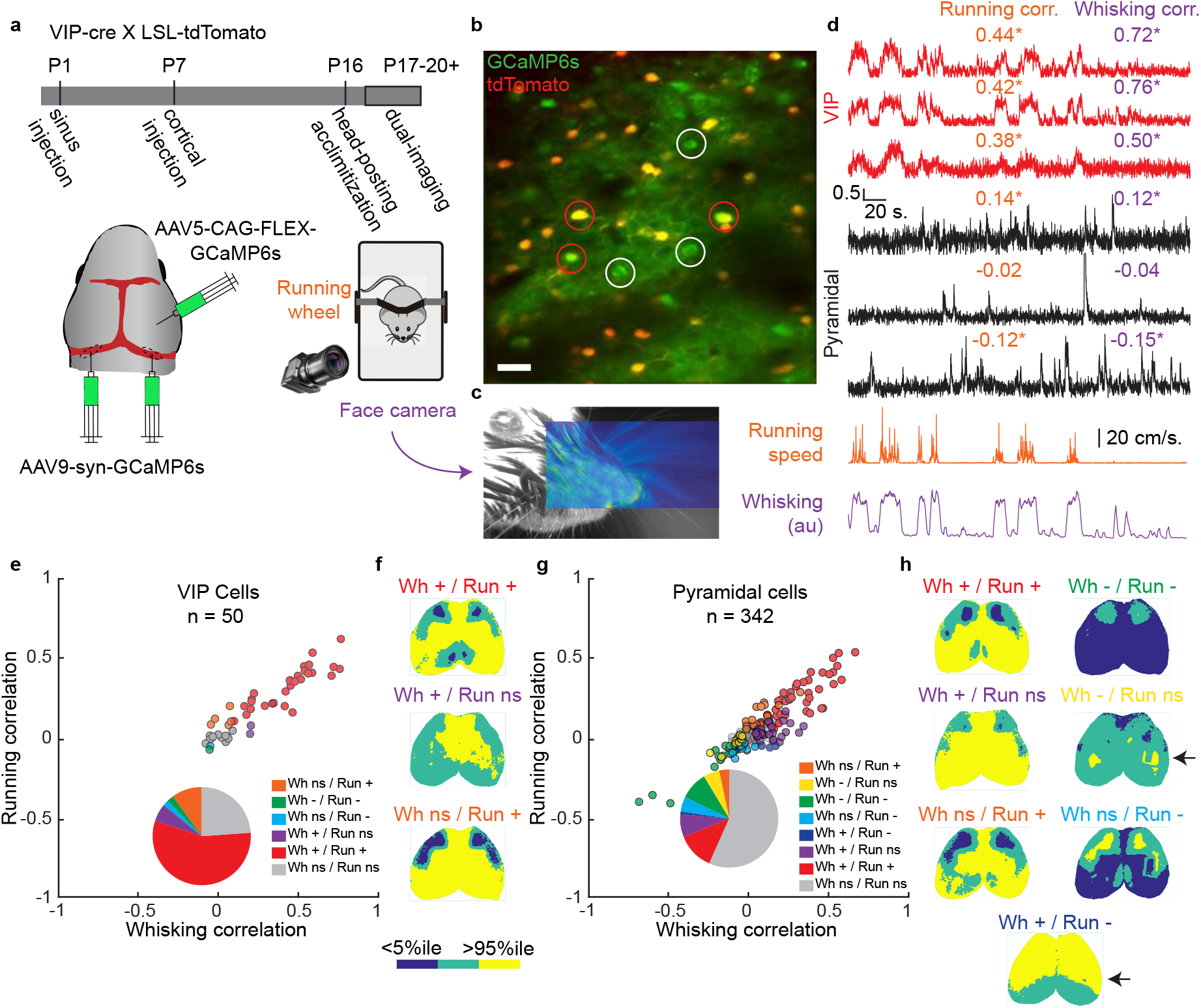
VIP-INs and pyramidal cells are linked to cortical networks associated with arousal state. **a**, Schematic indicating experimental timeline. **b**, Mean two-photon image from simultaneously acquired data showing GCaMP6s expression in tdTomato-positive (VIP-INs, examples circled in red) and -negative (presumptive pyramidal neurons, examples circled in white) cells. Scale bar is 20 μm. **c**, Example facial videography frame overlaid with whisker motion energy heat map. Warmer colors indicate higher mean pixel motion energy. **d**, ΔF/F traces from VIP cells (red) and pyramidal cells (black) with running speed (orange) and whisker motion energy (purple). Correlation of ΔF/F with running and whisking is listed above each trace, with asterisks indicating significant values (p < 0.01). **e**, Correlation with whisking and running for all analyzed VIP-INs. Colored points indicate cells with significant correlations. Inset pie chart shows distribution of significant correlations for VIP-INs. **f**, Significance maps for exemplar cells from the three groups of VIP-INs (groups with only one cell were excluded) with significant correlation to running and/or whisking. **g**, Same as e, but for pyramidal cells. **h**, Same as f, but for pyramidal cells, and with seven groups.

Of the 50 VIP-INs recorded, most were modulated by running (33/50), whisking (31/50), or both (28/50) (Figure 5e), and their CCNs were grossly similar, with significant posterior (often excluding retrosplenial cortex) activation and lateral frontal deactivation coincident with individual VIP-IN activity (exemplars shown in Figure 5f). Pyramidal cell response properties and CCNs were considerably more heterogeneous, with many cells either positively or negatively modulated by whisking and running (Figure 5g). Significant CCNs for pyramidal cells that were positively modulated by running and/or whisking were similar to those for VIP cells, while networks for cells negatively modulated by running had inverse patterns. Interestingly, though, cells that were negatively modulated only by whisking were co-active almost exclusively with bilateral S1, and cells that were positively modulated by whisking but negatively modulated by running were broadly co-active with the entire anterior cortex. The activity patterns of these latter two populations (arrows point to exemplars in Figure 5h), which are exclusive to pyramidal cells in our sample, suggest a dissociation in the cortex-wide network associations of cells that are modulated by running versus whisking.

## Discussion

The ability to relate the activity of individual, genetically-defined neurons to ongoing dynamics in distal cortical areas creates new opportunities to assess how different populations of cells participate in functional networks. We have presented a novel approach designed for this purpose, centered on a system for simultaneous two-photon cellular-resolution and whole-cortex widefield calcium imaging. We further present analytical approaches that demonstrate the unique advantages of this system.

With this dual imaging system, we have discovered that pyramidal cells in juvenile S1 have a greater diversity of cortex-wide network associations (CCNs) than has been shown previously for adults^14^ and that could be predicted from anatomical tracing studies^6,24,36^. In addition, we have found that VIP-INs, comprising a sparse yet critically important population of cells, are recruited as part of a common arousal network across medial and posterior cortex. In contrast, neighboring pyramidal cells have more diverse cortical network associations across behavioral state.

Importantly, these initial analyses relied on spontaneous activity in awake mice. However, our methods could easily be applied during sensory stimulation, in behavioral paradigms, and across a variety of disease models to investigate the structure and plasticity of cortical network architecture. For example, layer 2/3 neurons in S1 respond sparsely to whisker stimulation despite their necessity for whisker-guided behavior^37^. Using data from the same mice described in experiments here, we are currently assessing whether the activity of distal cortical areas gates the sparse coding of individual cells and whether there exist cortex-wide predictors of cellular response probability.

These imaging methods are enabled by the recent development of transgenic animals expressing bright, genetically-encoded calcium and structural indicators^18,29^. Recent advances in the development of similarly bright voltage^38^ and other functional indicators^39,40^ suggest the possibility of establishing the structure of cortical networks based on both electrophysiological and biochemical markers. Indeed, our transverse sinus injection technique for whole-brain gene delivery, along with other established methods for use in adult mice^32^, dramatically accelerates the development of novel molecular technologies without the need to generate costly transgenic strains.

Future methodological developments in our group are focused on improving the depth penetration and axial resolution of two-photon imaging through a prism through the use of red-shifted indicators and implementation of adaptive optics for correction of optical aberrations^41^. We are also adapting our current approach to include optogenetic activation and suppression of targeted cell types and cortical regions to assess the causal directionality of functional connections identified with our analyses.

Finally, these studies have been carried out in the context of a larger collaboration to relate neural activity across a broad range of spatiotemporal scales. We have also established an approach for carrying out mesoscopic calcium imaging simultaneously with functional magnetic resonance imaging (fMRI)^16^. This method will enable us to link the activity of targeted cells to brain-wide activity measured by blood oxygenation level-dependent (BOLD) signaling in mice as well as relating such activity to analogous fMRI studies in humans. Overall, we believe that the technical and conceptual union of these varied techniques will present powerful opportunities to drive novel understanding into the dynamic, functional architecture of networks in the mammalian nervous system.

## Methods

### Dual-imaging microscope design

Our dual-imaging microscope is composed largely of commercially-available components or items that can fabricated in a basic metalworking shop. The mesoscope is a Zeiss Axiozoom V.16 coupled to a PlanNeoFluar Z 1x, 0.25 NA objective with a 56 mm working distance. The mesoscope is mounted on a motorized Z-axis (Fluar Illuminator Z mot, Zeiss) with a manual X-axis dovetail slider (DTS50, Thorlabs) for precise positioning along two axes. Both the mesoscope and Z-axis motor are controlled remotely (EMS3 and Sycop3, Zeiss). Epifluorescence excitation is performed using a 7-channel LED driver (SpectraX, Lumencor) mated to the mesoscope by a 3 meter liquid light guide (Lumencor). The cyan LED channel is filtered using a ET470/20x filter (Chroma) and the violet LED channel is filtered using a ET395/25x filter (Chroma). LED illumination is reflected onto the imaging plane using a FT495 (HE) dichroic mirror and epifluorescence emissions are filtered with a BP525/50 (HE) filter (38 HE, Zeiss) and recorded using a sCMOS camera (pco.edge 4.2) with 512×500 resolution after 4×4 pixel binning. Images are acquired by a computer running Camware software (pco). The intensity of the LED excitation at the imaging plane is between 0.027 mW/mm^2^ and 0.106 mW/mm^2^ calibrated for each experiment depending on the brightness of the indicator.

The two-photon microscope is a Movable Objective Microscope (MOM) with the Janelia wide-path design and galvo-resonant scanner (Sutter Instruments) that has been customized for optimal imaging in a horizontal configuration. Specifically, one of the PMTs was re-positioned 180 degrees from its original position and springs were added to reduce strain on the micromanipulator. Two-photon excitation is performed using a Ti:Sapphire laser (MaiTai eHP DeepSee, Spectra-Physics) with built-in dispersion compensation. Laser intensity into the microscope is controlled using a Pockels cell (Conoptics) for rapid modulation and the laser is expanded and collimated with a 1.25x Galilean beam expander (A254B-100 and A254B+125, Thorlabs). The laser is focused on the brain using an ultra-long-working distance objective with a 20mm WD and 0.40NA (M Plan Apo NIR 20x, Mitutoyo) that was selected for high transmission of visible and near-IR light. We also tested other objectives and achieved similar results. Fluorescent emitted light is reflected into the collection path by a FF735Di-02 dichroic mirror (Semrock), filtered with a ET500lp long pass filter (Chroma) and then split by a T565lpxr dichroic mirror (Chroma) into two GaAsP PMTs (H10770PA-40, Hamamatsu) with ET525/50m-2p (Chroma) and ET605/70m-2p (Chroma) filters for detection of green and red photons, respectively. The combination of excitation, dichroic, and emission filters described here results in a net OD of >15 in between the cyan excitation illumination from the LED and the green emissions detected by the PMTs, which is sufficient to prevent contamination of the images formed by the two-photon microscope. The two-photon microscope is controlled using ScanImage 2017 (Vidrio Technologies) and images are acquired at 512×512 resolution with or without bi-directional scanning (see below).

### Synchronization of imaging modalities

For simultaneous widefield and two-photon imaging where the same emitted photons are being collected by the two modalities, we interleave frame acquisitions using a Master-8 (AMPI) stimulator to coordinate the timing and a Power3 DAQ and Spike2 software (Cambridge Electronic Design) to record all timing signals. Each “simultaneous” acquisition begins with a 33 – 66 millisecond two-photon acquisition (depending on whether bi-directional scanning is employed), followed by the simultaneous triggering of cyan LED excitation for 30 milliseconds and sCMOS camera frame acquisition for 20 milliseconds (the difference is due to the 10 milliseconds necessary for the rolling shutter of the camera to open) followed, in some experiments, by the simultaneous triggering of violet LED excitation for 30 milliseconds and a second sCMOS camera frame acquisition for 20 milliseconds. The sequence would then end with a 10 millisecond pause without any excitation or image acquisition, which was necessary to allow the PMTs to recover from the epifluorescence emissions excited by the LED. Data reported in Figures 2 and 3 were acquired without violet illuminated frames and bi-directional scanning and data reported in Figure 5 were acquired with violet illuminated frames and bi-directional scanning.

### Surgical preparation and imaging set up

All surgeries were performed in accordance with the regulations set by the Yale University Institutional Animal Care and Use Committee and in accordance with NIH guidelines. Mice were anesthetized using 1-2% isoflurane and were maintained at 37°C using an electric heating pad for the duration of the surgery. Meloxicam (0.3 mg/kg) was administered IP and topical lidocaine was applied to the scalp, which was then cleaned with povidone-iodine solution. The skin and fascia layers above the skull were removed to expose the entire dorsal surface of the skull from the posterior edge of the nasal bone to the middle of the interparietal bone, and laterally to the temporal muscles. The skull was thoroughly cleaned with saline and care was taken to not let the skull dry out. The edges of the skin incision were secured using cyanoacrylate (Vetbond, 3M). A custom headpost (see Figure 1a, inset) was secured to the interparietal bone and along the lateral edge of the right parietal bone, first with Vetbond, and then with transparent dental cement (Metabond, Parkell), and a thin layer of dental cement was applied to the entire exposed skull. In some animals, a metal “visor” was also secured to the nasal bone to prevent light from the wide-field microscope from getting in the animal’s eyes (although this could also be achieved by limiting the spread of light using a built-in aperture at the illumination port of the widefield microscope). Once the dental cement dried, it was covered with a thin layer of clear nail polish or cyanoacrylate (Maxi-Cure, Bob Smith Industries). The combination of the dental cement and cyanoacrylate dramatically increases the transparency of the skull.

For all imaging experiments described here, the mice were then allowed to recover for at least 3 hours in a heated recovery chamber and widefield imaging was performed to record spontaneous activity, as well as activity evoked by deflection of the left whiskers (10 Hz stimulation for 1 second every 40 seconds or a single 100 millisecond deflection every 10 seconds) using a piezo bender (PL112, Physik Instrumente), the latter of which was used to map the location of the right barrel cortex.

On the day following widefield imaging (or on the same day in some cases when the headpost implantation surgery occurred on a previous day – two surgeries were never conducted on the same day) the mice were then re-anesthetized and meloxicam (0.3 mg/kg) was administered IP, and a 2 mm square craniotomy was performed over the mapped location of the right barrel cortex and a 2 mm square BK7 glass microprism (Tower Optical) was lowered directly onto the surface of the dura and the edges were secured to the skull using a viscous cyanoacrylate (Gel Control, Loctite). In one mouse, a 1.5mm square craniotomy and glass microprism was used, and in two other mice, a glass coverslip was placed over the surface of the dura and the microprism was glued to the coverslip. Mice were then allowed to recover for at least 3 hours and either imaged on the same day as surgery or returned to their home cages for imaging on subsequent days. Juvenile mice pre-weaning were rubbed with bedding prior to being reintroduced into their litters to facilitate the dams’ continued care of the young mice.

For optimal two-photon imaging (minimizing reflections and aberrations), the front face of the microprism needs to be orthogonal to the long axis of the two-photon objective. Therefore, we iteratively adjusted the horizontal angle of the prism face by rotating the stage to which the animal is head-fixed and the vertical angle of the objective by rotating the microscope head. This procedure was performed until reflections off the front face of the prism were minimized, as assessed by monitoring the dim autofluorescence of the prism face as it moves through the imaging plane.

Using these procedures, we could typically image GCaMP6 up to 400 µm deep using laser powers at the front aperture of the objective of up to 300 mW. However, the power at the surface of the brain is likely significantly less as we did not observe photodamage or photobleaching arising from the two-photon imaging.

This imaging preparation is stable over days to weeks if care is taken to clean and protect the implanted prism from debris in the animal’s home cage (or damage due to grooming behaviors). At the termination of an experiment, the microprisms could be recovered and cleaned with acetone, followed by 2M HCl, followed by 100% methanol, in which they could also be stored until a subsequent implantation.

### Animal subjects

For dual-imaging experiments, we used either Slc17a7-cre/Camk2a-tTA/TITL-GCaMP6s^29,42^ (The Jackson Laboratory strains 023527, 024108) or VIP-cre/LSL-tdTomato^43,44^ (The Jackson Laboratory strains 010908, 007909) mice. Mice were housed on a 12-hour light/dark cycle with food and water available *ad libitum*. For histological validation of sinus injections, we used C57BL6/J mice.

### Transverse sinus injections

Postnatal-day-1 pups were taken out of their home cage with their entire litter and rested on a warm pad. Pups were anesthetized using hypothermia which was induced by laying them on ice for 2-3 minutes. They were then maintained on a cold metal plate for the duration of the procedure. A light microscope was used to visualize the transverse sinuses (located on the dorsal surface of the mouse head between the parietal and interparietal bones). Fine scissors were used to make two small cuts (∼2 mm) in the skin above each transverse sinus. To inject the virus, we used capillary glass tubes, pulled using a P-97 pipette puller (Sutter Instruments) to produce fine tips with high resistance. The sharp pipettes were filled with mineral oil and attached to a Nanoject III. Next, most of the mineral oil was pushed out of the pipette using the Nanoject, and vector solution was drawn into the pipette. For accurate movement of the Nanojet, we used a MP-285 micromanipulator (Sutter Instruments). The pipette is gently lowered into the sinus until the tip of the pipette has broken through the sinus. The pipette tip was then raised back until it was 300-400 µm below the surface of the sinus. Next, we injected 4 µL of AAV9-Syn-GCaMP6s (Addgene) with a titer of ∼3×10^13^ vg/mL. Injections were performed at a rate of 20 nL/second. A correct injection was verified by blanching of the sinus. After the injection, the skin was folded back, and a small amount of Vetbond glue was applied to the cut. The pup was then returned to the warm pad. After the entire litter was injected, the pups were returned to their home cage and gently rubbed with bedding to prevent rejection by the mother. At 13 or 20 days post-injection, juvenile mice were perfused with PBS and 4% paraformaldehyde (PFA) solution and brains were extracted, immersion fixed in PFA overnight, and rinsed with PBS.

### Histological processing and immunohistochemistry

Juvenile brains were cut into 150 µm coronal or sagittal slices using a vibratome (Leica). Slices were transferred into 0.04% Triton solution, then blocked overnight with 10% goat serum at 4°C. After blocking, primary antibodies were diluted in the blocking solution (1:500), and slices were incubated in the primary antibody solution for 5 days at 4°C. The primary antibodies used were rabbit anti-GFP conjugated to AF488 (Millipore) and mouse anti-NeuN (Millipore). Slices were then washed three times with PBS, and incubated in secondary antibody diluted in blocking solution (1:500) overnight at 4°C. The secondary antibody used was goat anti-mouse AF555 (Millipore). Slices were washed three times with PBS, incubated in DAPI diluted in PBS (1:1000) for 15 min, washed three times with PBS and mounted on glass slides using Fluoromount G. Sagittal slice images were captured using a Zeiss Apotome microscope, and coronal slices used for quantification were imaged using a laser scanning confocal microscope (LSM 800, Zeiss) to determine co-localization between anti-GFP and anti-NeuN signals. Signal quantification was done using ImageJ software. Briefly, regions of interest (ROIs), i.e. cortex and thalamus, were selected, images were binarized, and the number of GCaMP6s+ and NeuN+ cells were counted manually.

### Injections for VIP experiments

For experiments in which the activity of individual VIP cells was measured simultaneously with mesoscopic pyramidal cell activity across the cortical mantle, we performed transverse sinus injections at P1, as described above. Sinus injection of AAV9-Syn-GCaMP6s preferentially labels pyramidal cells (data not shown). Therefore, to enhance the expression of GCaMP6s within VIP cells in S1, we also performed cortical injection of AAV5-CAG-Flex-GCaMP6s (Penn Vector Core) at P6. Between 400 – 600 nL of virus was injected using a Nanoject III. To perform these injections, a sharp glass pipette was cut to a 10 µm tip diameter and beveled at a 45° angle. Mice were anesthetized using 1-2% isoflurane and maintained at 37° on a water-recirculating heating pad. Meloxicam (0.3 mg/kg) was administered IP and lidocaine (0.5%) was administered locally SQ for analgesia. An incision was made to expose lambda and the pipette tip was moved with the micromanipulator to +1.8 mm anterior, +2.0 mm lateral from lambda, just above the surface of the skull. The skull was scraped with a scalpel blade at that location to thin it and then the pipette tip was then slowly lowered until it pierced through, typically within 800 µm from the skull surface. The tip was then slowly withdrawn to a depth of 400 µm below the surface of the skull and the brain was allowed to settle for 5 minutes prior to injecting the virus at a rate of 2 nL/second. 5 minutes after completion of injection, the pipette was withdrawn and the incision was closed with Vetbond. The pup was allowed to recover on a warm heating pad. Prior to returning injected mice to their home cage, they were mixed with all of their littermates (injected and uninjected) and rubbed with bedding to prevent rejection by the dam.

### Behavioral monitoring

To perform simultaneous widefield and two-photon imaging in awake, behaving mice, we head-fixed the mice such that they could run freely on a cylindrical running wheel, and the angular position of the wheel was recorded as described previously^22^. To assess the behavioral state of the mice, we performed videography of the face, including the whiskers and pupil, using a miniature CMOS camera (Flea3, FLIR) with a variable zoom lens (13VM20100AS, Tamron). The face was illuminated with an 850 nm NIR LED array. Face image acquisition was time-locked to the onset of the two-photon acquisition to ensure constant luminance across frames. Therefore, for the experiments described here, facial videography occurred at 9.55 Hz (experiments described in Figures 2 and 3) or 9.15 Hz (experiments described in Figure 5). During all recordings described here, no additional sensory stimulation was provided to the mice.

### Analysis methods

#### Two-photon data pre-processing

Two-photon data was motion corrected for x-y displacements by rigid body transformation using the moco toolbox^19^ in ImageJ (NIH). For data acquired from VIP-cre/LSL-tdTomato mice, motion correction was performed on the tdTomato frames and the calculated offsets were applied to the GCaMP frames. Motion-corrected frames were then top-hat filtered across time to compensate for whole frame changes in brightness and ROIs were manually selected and neuropil signal was removed from each ROI’s fluorescence signal as described elsewhere^9,18^. ΔF/F was calculated for each cell using the 10^th^ percentile as the baseline. In some cases of cells expressing both GCaMP6 and tdTomato, we observed bleaching in the green fluorescence intensity over a 20 minute dual-imaging. To compensate, we fit the baseline of each cell’s ΔF/F trace with a second-order exponential and removed the decay component. We use skewness as a proxy measure for cell activity, calculating skewness for each cell’s ΔF/F trace and excluding all cells with a value less than 0.5. For all experiments, we confirmed by visual inspection that this threshold results in a very low false positive rate for both pyramidal and VIP cells.

#### Widefield data pre-processing

Widefield data was rotated to align the anterior posterior axis with vertical and then a mask was super-imposed to remove pixels outside of cortex as well as over the superior sagittal sinus. Slow drifts in the baseline fluorescence of each pixel were removed using top-hat filtering across time (300 frame filter object). For data acquired without interleaved violet illuminated frames, ΔF/F for each pixel was calculated using the top-hat filtered traces and setting the baseline to the 10^th^ percentile value for each pixel across time. To remove changes presumably attributable to motion or hemodynamic artefacts, we performed global signal regression, which zero-centers the data, and then spatially smoothed the data using a Gaussian filter (σ = 2). For data acquired with interleaved violet illuminated frames, we performed pixel-wise regression of the (top-hat filtered) violet illuminated fluorescence values from the cyan illuminated fluorescence values. GCaMP fluorescence emissions under violet illumination are largely calcium-independent^18^. Therefore, fluctuations in fluorescence values should be largely attributable to (sub-pixel or z-axis) brain motion or hemodynamic artefacts. ΔF/F for each pixel was then calculated using the mean fluorescence value for each pixel across time and each frame was spatially smoothed using a Gaussian filter (σ = 2).

#### Behavioral state measures pre-processing

To calculate running speed, the angular position of the wheel (recorded at 5000 Hz) was down-sampled by a factor of 100 and converted to real displacement. To quantify whisking, we cropped the facial videos to only include the area corresponding to the whisker and calculated motion energy as the frame-to-frame change in intensity of each pixel. In addition to the displacement of the whiskers, this measure also reflects movements of the mouse’s nose.

#### Spike-weighted activity map calculation

Spike-weighted activity maps were calculated by taking the dot product of a matrix of ΔF/F values for all widefield pixels over time and each cell’s normalized spike probability vector. The single cell spike probability vectors were calculated using constrained-foopsi^27^ by the integral of each cell’s spike probability over time. This method has two advantages – the first is to increase the weighting of the onset of calcium transients to more accurately approximate the timing of cell spiking and the second is to center all of the derived network map pixel values to the mean of their activity over time. This facilitates easy comparison of CCNs within animals. To determine which cortical areas are significantly activated or deactivated coincident with the activity of individual cells, we generated a null distribution of CCNs for each cell by randomly circularly shifting the timing of cell spike probability relative to pixel ΔF/F. This method is necessary due to the statistical dependence of successive timepoints in a cell’s spike probability vector. All pixels below the 5^th^ percentile or above the 95^th^ percentile of the distribution were classified as significantly deactivated or activated, respectively. For all significance maps shown, blue pixels are referred to as “deactivated” and yellow pixels are referred to as “activated.”

We have compared our method to similar methods from other groups (calculating the correlation of cell activity and pixel activity^15^ or averaging the activity of time points when the cell is estimated to have fired an action potential^14^) and found that results were consistent across methods.

#### Parcellation of widefield imaging data

For the anatomical parcellation, we used a parcellation based off of the Allen Common Coordinate Framework version 3 (CCFv3). We combined some small parcels, such as the higher-order visual areas (8 defined areas combined into 2 parcels), to create a 16-node-per-hemisphere parcellation, which we aligned to each mouse using the superior sagittal sinus, transverse sinuses, inferior cerebral veins, and the stimulation evoked positions of barrel and auditory cortex. To compensate for differences in the angle of the skull between mice, we scaled the two hemispheres independently.

For the individualized functional parcellation, we applied a multi-graph k-way spectral clustering algorithm, as is described in greater detail elsewhere^16^. For each mouse, the parcellation was determined using widefield-only imaging data collected prior to microprism implantation. In this way, we avoid overfitting of the activity within each session. To apply the parcellation to the data collected during subsequent dual-imaging trials, we calculated the transformation to register the dual-imaging acquisitions to the widefield acquisition using an automated intensity-based registration and then applied the same transformation to the functional parcellation. For all animals, the transformation was rigid, except for one animal for which the parcellation was also scaled. We set the number of parcels to 16 per hemisphere in order to match the anatomical parcellation. The functional parcellations of the two hemispheres were obtained separately, thus there is no guarantee that the parcels from the two hemispheres look symmetric. Moreover, the difference in the vasculature had a notable impact on the parcellation results. In spite of all this, the functional parcellation shows a great degree of symmetry across a range of number of parcels from N=2 to N = 20 per hemisphere.

We observed that, in general, the CCN and significance maps are better aligned with the functional parcellation. Particularly, the functionally defined parcels have a good overlap with the activated/deactivated regions in each cell’s significance map, and the centers of the activated/deactivated regions align with the centers of functional parcels. We further quantified the alignment between the significance maps and the parcellations using an information theory measure called conditional entropy H(Y/X). Y represents the significance map and X represents the parcellation (anatomical or functional). If the significance map aligns with the parcellation, the information contained in X and Y are similar, thus the conditional entropy will be low. If Y is completely determined by X, then H(Y/X) = 0. On the contrary, if Y is independent of X, H(Y/X) will be high. Two conditional entropy values were calculated for each neuron (same significance map), one based on the functional parcellation and the other based on the anatomical parcellation.

#### Clustering of spike-weighted activity maps

After obtaining the significance map for each selected neuron, we calculated the ratio between the total number of significant (activated or deactivated) pixels within a parcel and the total number of pixels of the same parcel. This ratio can be viewed as a measure of average activation within a parcel. Accordingly, we created a feature vector of size 1 by 64 for each neuron, where the first half of the vector corresponds to the average activation from all 32 parcels and the second half corresponds to the average deactivation. To find the representative mesoscopic patterns of activation related to neuronal activity, we applied the spectral clustering algorithm (same algorithm used to generate the functional parcellation) to divide the neurons into three groups based on the feature vectors. For each of the three groups, we selected one neuron that is the most similar to all other neurons in the same group, and used its significance map as the exemplar of the group. For graphical purposes, we combined the positive and negative feature vectors into a single 1 by 32 vector for each neuron with values from −1 (all pixels significantly deactivated) to 1 (all pixels significantly activated), which we label the “activity index.”

#### Analysis of cellular response to running and whisking

We calculated the Pearson’s correlation of each cell’s ΔF/F trace with simultaneously acquired running velocity and whisker motion energy traces. We determined the significance of this correlation by performing 1000 random circular shifts of the timing of running or whisking relative to cell activity and setting significance as less than the 1^st^ percentile or greater than the 99^th^ percentile. Exemplar cells (in Figure 5f,h) were chosen by calculating the centroid of the whisking and running correlation values for all cells in each group and selecting the cell with the shortest Euclidean distance from the centroid.

## Code availability

All data and code used for these analyses will be made available upon reasonable request.

## Supplementary Figures

**Supplementary Figure 1.**
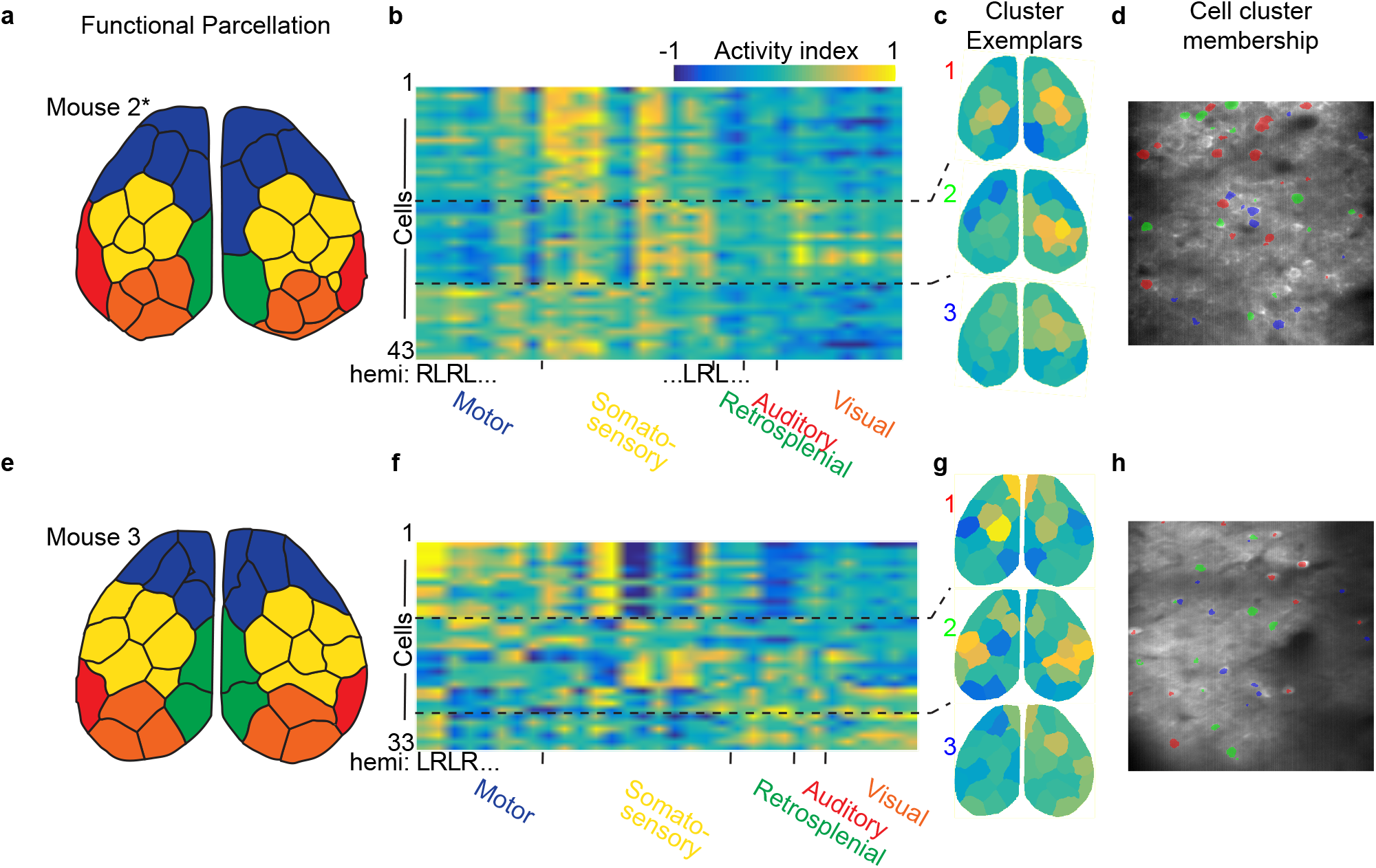
Cortex-wide CCNs of pyramidal cells identified with dual micro- and mesoscopic imaging. **a**, Functional parcellation of data from an example mouse. For this case, the parcellation algorithm calculated 15 parcels for the left hemisphere. **b**, Activity index calculated for all significance maps using the functional parcellation, ordered by cluster membership. **c**, Exemplars from the 3 clusters in b. **d**, Two-photon field-of-view with individual cells colored by cluster membership. **e-f**, Same as a-d, but for a different mouse.

